# Re-examination of the taxonomic status of the Antarctic *Pseudomonas syringae* Lz4W isolate and proposal to rename it as a novel species *Pseudomonas cryophila* sp. nov

**DOI:** 10.1101/2024.04.22.590509

**Authors:** Malay K Ray, Apuratha Pandiyan, Bhubanananda Sahu

## Abstract

A taxonomic re-evaluation of the Antarctic psychrotrophic bacterium *Pseudomonas syringae* Lz4W was performed in the light of its available genome sequence and due to a revision in the key phenotypic characteristics that are in conflict with the “*syringae*” group of Pseudomonads. A 16S rRNA gene sequence based phylogenetic analysis suggested that Lz4W^T^ strain belongs to “*fragi*” cluster of *Pseudomonas* species, with closest similarity (99.72%) to the type strain *P. deceptionensis* M1^T^. However, *in silico* analysis of the Lz4W genome sequence using SpecI (species identification tools), ANI (average nucleotide identity), and GBDP (Genome Blast Distance phylogeny) methods suggest that Lz4W^T^ strain cannot be delineated with any of the type strains of “*fragi*” cluster of species. Based on predictive low DNA-DNA hybridization value (<29.9%) and differences in phenotypic features with the related species we suggest that Lz4W^T^ is a novel species under the *Pseudomonas* genus, and we propose that the strain be named as *Pseudomonas cryophila* sp. nov. The type strain is Lz4W^T^ (=CFBP 8403^T^ =KCTC 42933^T^ =LMG 29591^T^ =MTCC 673^T^).

---

Gram-negative *Pseudomonas* strains are ubiquitous in nature. Their ability to survive in different extreme conditions makes them ideal for understanding microbial adaptation and evolution. We have been using an Antarctic isolate, formerly classified as *Pseudomonas syringae* Lz4W (Shivaji *et al*., 1989) for investigating the molecular mechanism of cold-adaptation for many years (Ray *et al*., 1998; Ray, 2006). This bacterial strain was isolated from the soil samples collected in and around the lake Zub of Schirmacher Oasis in east Antarctica during an Indian expedition (1984-1985), and identified as *P. syringae* Lz4W (Lz stands for lake Zub, and 4W for white colony number 4). The Lz4W^T^ is capable of growing at a temperature range of 0 to 30°C, under the laboratory conditions. In the Antarctic bacterial medium (ABM), which contains half the nutrients of LB (Luria-Bertani) medium but without sodium chloride, Lz4W^T^ grows with a generation time of ∼4.5 hrs at 4° C, but shows ∼1.5 hrs generation time at 22-24°C. Morphologically, Lz4W^T^ undergoes cell size variation, being reduced at low temperature (Regha *et al*, 2005) or in minimal growth medium (Sahu & Ray 2008). Biochemical and genetic studies with Lz4W^T^ have established various cold-adaptive mechanisms that allow the Antarctic bacterium to sense and grow at low temperature (Ray *et al*., 1994a, 1994b, 1994c, 1999; Uma *et al*., 1999; Jagatap & Ray 1999; Janiyani & Ray, 2002; Seshukumar *et al*., 2002; Regha *et al*., 2005; Purusharth *et al*., 2005, 2007; Satapathy *et al*., 2008; Singh *et al*., 2009; Pavankumar *et al*., 2010; Sulthana *et al*., 2011; Jagannadham & Chowdhury, 2012; Sinha *et al*, 2013; Kulkarni *et al*., 2014). However, the taxonomic identification of Lz4W^T^ as *P. syringae* species has remained ambiguous, as few key biochemical features of the strain reported earlier (Shivaji *et al*., 1989) were found incorrect, and hence identification of this strain as *P. syringae* species was untenable (Cindy Morris, personal communication). A global study on the *P. syringae* species diversity indicated that the Lz4W^T^ strain lies outside the *P. syringae* complex and it is closer to *P. protegens* Pf-5 species (Berg *et al.,* 2014). Thus, it was becoming clear that Lz4W^T^ should be classified as a separate species of *Pseudomonas*, and cold-adaptive features of the strain should be referred to under the new species and NOT under the “*P. syringae*” species. This will facilitate communication between the people using this strain for research and remove any confusion in scientific literatures (Moreno & Rojo, 2014; Ait-Bara & Carpousis, 2015) for its present classification under *P. syringae* species. Towards this goal, we took a genome based approach to re-examine species status of the Lz4W^T^ isolate. Here we report SpecI (Mende *et al*., 2013), JSpecies (Richter & Rossello-Mora, 2009), and GBDP (Meier-Kolthoff *et al*., 2013) based analysis of Lz4W^T^ genome sequence (accession no. AOGS00000000.1), along with the 16S rRNA gene and multilocus sequence analysis (MLSA) based phylogeny. Taking the revised phenotypic characteristics and genome sequence based species delineation into account, we suggest that the Antarctic Lz4W isolate be classified as *Pseudomonas cryophila* Lz4W^T^ sp. nov.

First, we put in record few key phenotypic characteristics that were reported earlier in description of Lz4W^T^ (Shivaji *et al*., 1989) and were found incorrect in the reassessment of these tests (Table 1). For example, Lz4W^T^ isolate was scored negative for the oxidase and arginine dihydrolase tests, which were later observed to be positive. The strain was also incapable of hydrolyzing esculin and negative for the induction of hypersensitive reaction in tobacco (Cindy E Morris, 2005, personal communication). We also performed several carbon source and amino acids utilization properties of the strain Lz4W^T^, by using the defined minimal growth medium (MM_Lz_: 8.45 mM Na_2_HPO_4_, 4.41 mM KH_2_PO_4_, 3.79 mM (NH_4_)SO_4_, 1.0 mM MgSO_4_, 0.01% of valine and isoleucine and desired carbon source) that was developed to study carbon catabolite repression control in the bacterium (Sahu & Ray, 2008). These test results (Table 2) were also in variance with the reported abilities of Lz4W^T^ strain for the carbon source and amino acids utilization (Shivaji *et al*., 1989). Lz4W^T^ strain did not utilize sucrose and mannitol as carbon source, as reported earlier. It utilized histidine, asparagine, serine, phenylalanine, tyrosine and glycine as carbon and nitrogen sources when added to minimal medium (MM_Lz_). The growth on glycine, and threonine was relatively poor. It is to be noted that the observed variations in carbon and nitrogen source utilization might be related to the use of soil extracts as supplement to growth medium by Shivaji *et al*. (1989). Here we have used MM_Lz_ containing defined mineral salts, and isoleucine and valine (0.01% each) for the carbon source and amino acids utilization tests. This is important as Lz4W^T^ was found to be a natural auxotroph for the amino acids isoleucine and valine, and could not be grown in standard M9 basal medium (Sahu & Ray, 2008). Additionally, we observed that Lz4W^T^ does not grow in 5.8% NaCl as reported earlier (Shivaji *et al*, 1989) and this is consistent with the finding that Lz4W^T^ grows best in low osmolar (∼45 mOsmol) medium (Sahu & Ray, 2008). We also note that the reported G+C content (64.4%) estimated from the T_m_ value of genomic DNA by Shivaji *et al*. (1989) is in variant with the actual G+C content (58.67%) that has been derived from the genome sequence of Lz4W^T^ (Pandiyan & Ray, 2013).

**Table 1.** Major phenotypic characteristics that differentiate Lz4W^T^ isolate from the “*syringae*” group of *Pseudomonas*. A ‘+’ and ‘−’ indicates positive and negative for the tes ts, respectively for the *P. syringae* CFBP1392^T^ type strain. The % values shown for tests-positive (+) reactions in bracket are from Berge *et al*. (2014) indicating concurrence of the reactions by different “*syringae*” strains grouped under the ‘*P. syringae* complex’ collected from different environments of the globe.

| Characteristics | Lz4W<sup>T</sup> isolate | “*syringae*” group |
| --- | --- | --- |
| Oxidase | + | - |
| Arginine dihydrolase | + | - |
| Esculin hydrolase | - | + (97%) |
| Hypersensitive reaction in Tobacco | - | + (73%) |
| Fluorescent pigment production in King’s B medium | - | + (98%) |
| Levan production | - | + (41%) |
| Sucrose utilization | - | + (73%) |

**Table 2.** A few revised and additional characteristics of carbon source and amino acid utilization by the *Pseudomonas* Lz4W^T^ isolate. A ‘+’ and ‘−’ indicates positive and negative for the tests, respectively. nd, not determined.

| **Carbon source utilization** | Revised in this study | Reported in Shivaji *et al* (1989) |
| --- | --- | --- |
| sucrose | - | + |
| mannitol | - | + |
| fumerate | + | nd |
| α-ketoglutarate | + | nd |
| D-glucuronate | + | nd |
| **Amino acid utilization** |  |  |
| L-methionine | - | + |
| L-histidine | + | - |
| L-asparagine | + | - |
| L-serine | + | - |
| L-Phenyl alanine | + | - |
| L-tyrosine | + | - |
| glycine | + (poor) | - |
| **Salt tolerance** |  |  |
| 5.8% NaCl | - | + |
| **G+C content** |  |  |
| GC % | 58.7 | 64.4 |

To re-examine the taxonomic status of Lz4W^T^ by 16S rRNA gene homology, we constructed a maximum-likelihood (ML) phylogenetic tree using the 16S rRNA gene sequences of *Pseudomonas* type strains that showed >97% sequence similarities to Lz4W^T^ (Fig. 1 & Table S1). The sequences were extracted from the Ez-Taxon server (Kim *et al*. 2012), aligned using MUSCLE with default parameters (Edgar 2004), and phylogenetic tree was constructed using MEGA 6 software (Tamura *et al*., 2013). Lz4W^T^ 16S rRNA gene displayed highest similarity with different type strains of the *P. fragi* subgroup of species, *P. deceptionensis* M1^T^ (99.72%), *P. endophytica* BSTT44^T^ (99.52%), *P. psychrophila* E-3^T^ (99.46%), *P. lundensis* ATCC49968^T^ (99.46%), and *P. fragi* ATCC4973^T^ (99.45%). This high degree of similarity led us to additionally perform MLSA using the concatenated sequences of 16S rRNA, *gyrB*, *rpoB* and *rpoD* genes of the related *Pseudomonas* species (Mulet *et al,* 2010). The MLSA based ML phylogenetic tree (Fig. 2) also placed Lz4W^T^ in the *P. fragi* cluster of species, but with 95.99, 95.86, 95.65, 93.6, and 92.97% similarity with the type strains of *P. psychrophila*, *P. fragi*, *P. deceptionensis*, *P. lundensis*, and *P. endophytica*, respectively. Interestingly, *P. endophytica* BSTT44^T^ which was found closest to Lz4W^T^ in the 16S rRNA gene analysis became closeted with *P. lundensis* in the MLSA based tree. Nonetheless, MLSA scores were below the species threshold (>97% similarity) suggesting that the Antarctic Lz4W^T^ strain might be a new species.

**Fig. 1.**
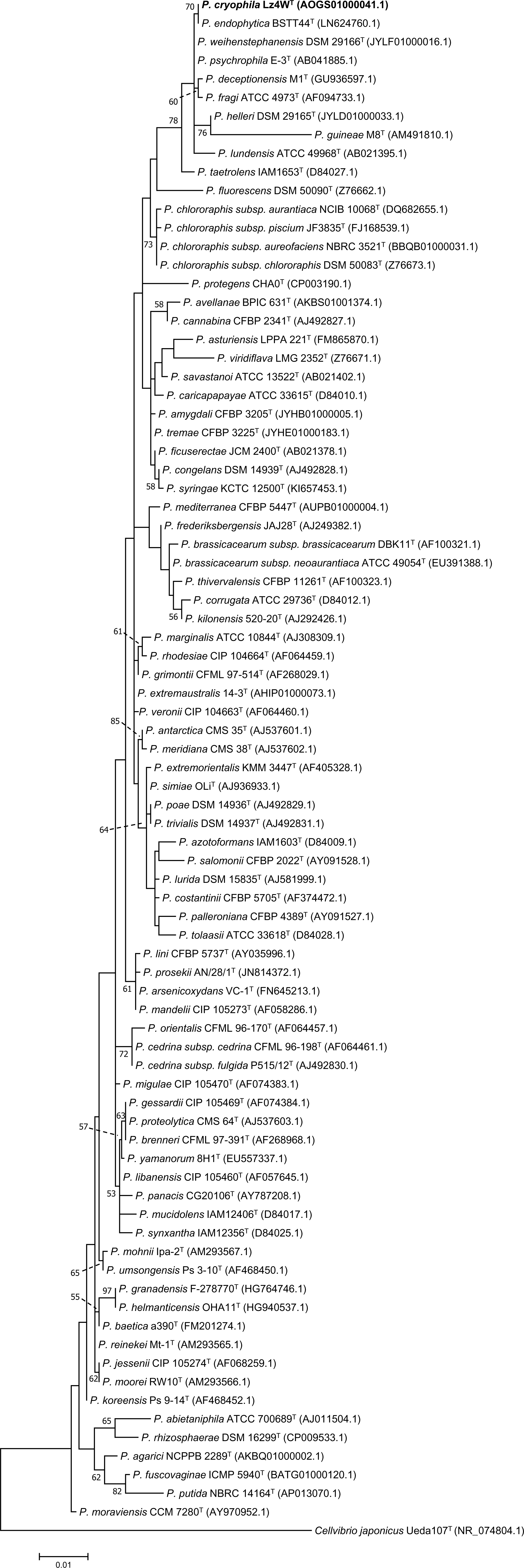
Maximum-likelyhood (ML) phylogenetic tree based on 16S rRNA sequence of the Lz4W^T^ and closely related type strains of the genus *Pseudomonas* showing >97% sequence similarity. *Cellvibrio japonicus* Ueda107^T^ was used as outgroup for the analysis. Bootstrap values >50% (based on 1000 replicates) are shown at branch points. GenBank accession numbers are indicated in brackets. Scale bar: 1 nucleotide substitution per 100 sites.

**Fig. 2.**
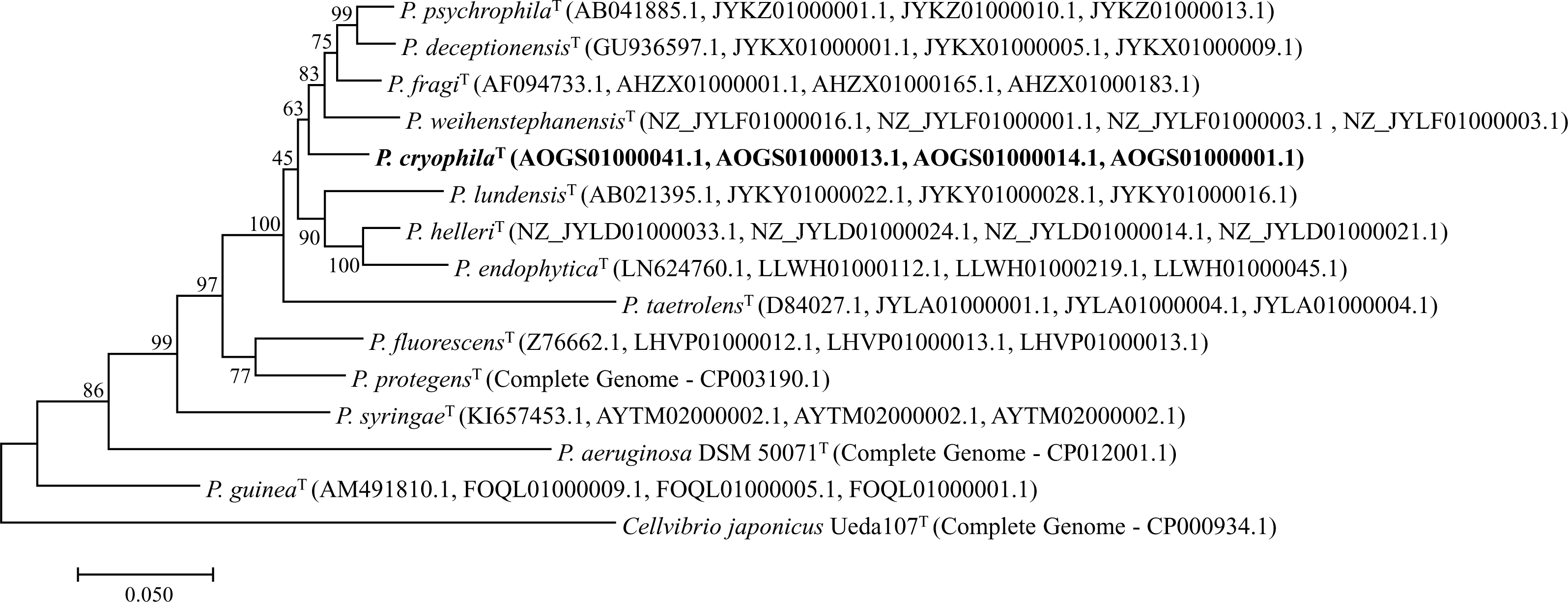
ML phylogentic tree based on concatenated sequence of 16S rRNA, *rpoB*, *rpoD*, and *gyrB* gene sequences of Lz4W^T^ and phylogenetically closely related type strains of selected species of *Pseudomonas* genus. *Cellvibrio japonicus*^T^ was used as outgroup. Bootstrap values >50% (based on 1000 replications) are shown next to the branches. Genbank accession numbers are indicated within brackets. Scale bar, 5 nucleotide substitution per 100 sites.

With the ease of genome sequencing methods and low cost, the numbers of sequenced genomes have increased exponentially, leading to the development of genome-sequence based several novel methods of classification of bacterial species (Konstantinidis & Tiedje, 2005). We first used SpecI (species identification tool) to assign the species identity of Lz4W strain. This web based analytical tool (Mende *et al*., 2013) uses the sequence data of organisms for which complete genome sequence is known. The species are assigned based on identity score of 40 universal, single copy phylogenetic marker genes (PMGs) that are present in all three domains of life. SpecI webserver (www.bork.embl.de/software/specI/) automatically extracts the 40 universal PMGs from a given genome and performs distance calculations to a database of the marker genes from ∼3500 genomes in the data base. The query genome is then assigned to a species if the PMGs are on average more than 96.5% identical to all genomes of the assigned species represented by one cluster. In this analysis, Lz4W^T^ was found to be closer to *P. fluorescens* Pf-01 strain, with an average identity score of 89.58%, much below the species cutoff values of 98.5% for assigning the Lz4W strain to *P. fluorescens* species. None of the draft genome sequences of the *P. fragi* group of species were available in database of the SpecI web server. We therefore extracted nucleotide sequence of the 40 PMGs (Table S2) from the draft genome sequences of *P. fragi* group of species that were available at the NCBI, and calculated the average percentage identity of each PMG by using default parameters of nucmer (MUMmer) program, except for *ftsY* gene (COG00552). The *ftsY* showed only partial alignment, and hence its sequences were extracted using BLAST and then a global alignment was performed using EMBOSS-NEEDLE (www. ebi.ac.uk). The SpecI identity scores (Table 3) shows that Lz4W^T^ exhibits highest average identity (94.49, 93.90, 93.60, 91.79, and 91.4 %) with the five type-strains *P. deceptionensins* M1^T^, *P. psychrophila* E-3^T^, *P. fragi* ATCC4973^T^, *P. lundensis* ATCC49968^T^, and *P. endophytica* BSTT44^T^ respectively, of the *P. fragi* species cluster. These values underscore that Lz4W^T^ is closely related to ‘*fragi*’ subcluster of species, but the identity scores are below the species cutoff value (98.5%) and hence Lz4W^T^ cannot be assigned to any of these species.

**Table 3.** SpecI analysis of genome identity for establishing taxonomic relationship of the Lz4W^T^ with closely related “*P. fragi* cluster” of type strains. DNA sequences of 40 PMGs (Phylogenetic Marker Genes) were extracted from the draft genome sequences of the indicated strains available at the NCBI site. The length-weighted average identity values for the 40 PMGs are shown at the bottom row of table. The species are: **1**, *P. deceptionensis* M1^T^; **2**, *P. psychrophila* E-3^T^; **3**, *P. fragi* ATCC4973^T^; **4**, *P. taetrolens* DSM21104^T^; **5**, *P. lundensis* DSM6252^T^; **6**, *P. endophytica* BSTT44^T^. The COG (Cluster of Orthologous Groups) numbers of 40 PMGs and their respective locus tags in the Lz4W^T^ genome have been indicated.

| COG # of 40 PMGs | Protein | Lz4W Genome Locus-Tag | % Identity with |  |  |  |  |  |
| --- | --- | --- | --- | --- | --- | --- | --- | --- |
|  |  |  | 1 | 2 | 3 | 4 | 5 | 6 |
| COG0012 | Predicted GTPase, probable translation factor | B195_04736 | 92.35 | 91.89 | 91.71 | 91.89 | 87.89 | 89.07 |
| COG0016 | Phenylalanine-tRNA synthetase, alpha subunit | B195_09269 | 93.81 | 93.31 | 93.51 | 93.31 | 88.4 | 91.54 |
| COG0018 | Arginyl-tRNA synthetase | B195_02588 | 90.39 | 90.16 | 89.42 | 90.16 | 88.32 | 86.45 |
| COG0048 | Ribosomal protein S12 | B195_13621 | 98.92 | 98.39 | 98.66 | 98.39 | 97.31 | 97.58 |
| COG0049 | Ribosomal protein S7 | B195_13616 | 97.66 | 97.45 | 97.03 | 97.45 | 96.6 | 94.90 |
| COG0052 | Ribosomal protein S2 | B195_05980 | 98.25 | 97.98 | 98.25 | 97.98 | 96.37 | 96.37 |
| COG0080 | Ribosomal protein L11 | B195_13651 | 96.06 | 95.6 | 95.83 | 95.6 | 95.6 | 95.14 |
| COG0081 | Ribosomal protein L1 | B195_13646 | 96.12 | 96.41 | 95.26 | 96.41 | 92.67 | 91.68 |
| COG0085 | DNA directed RNA Polymerase, beta subunit | B195_13631 | 95.7 | 95.75 | 96.15 | 95.75 | 92.19 | 92.86 |
| COG0087 | Ribosomal protein L3 | B195_13596 | 99.06 | 99.21 | 97.48 | 99.21 | 98.58 | 96.23 |
| COG0088 | Ribosomal protein L4 | B195_13591 | 97.01 | 96.85 | 96.68 | 96.85 | 96.85 | 96.52 |
| COG0090 | Ribosomal protein L2 | B195_13581 | 99.27 | 96.97 | 96.97 | 96.97 | 95.52 | 96.00 |
| COG0091 | Ribosomal protein L22 | B195_13571 | 99.7 | 99.1 | 99.1 | 99.1 | 99.1 | 98.80 |
| COG0092 | Ribosomal protein S3 | B195_13566 | 98.84 | 98.84 | 99.13 | 98.84 | 97.96 | 97.67 |
| COG0093 | Ribosomal protein L14 | B195_13546 | 100 | 100 | 100 | 100 | 98.1 | 98.10 |
| COG0094 | Ribosomal protein L5 | B195_13536 | 97.59 | 97.59 | 98.15 | 97.59 | 96.11 | 94.26 |
| COG0096 | Ribosomal protein S8 | B195_13526 | 98.73 | 99.75 | 98.98 | 99.75 | 97.71 | 95.17 |
| COG0097 | Ribosomal protein L6P/L9E | B195_13521 | 98.31 | 99.25 | 98.69 | 99.25 | 97.19 | 96.26 |
| COG0098 | Ribosomal protein S5 | B195_13511 | 96.61 | 96.41 | 96.61 | 96.41 | 97.01 | 95.21 |
| COG0099 | Ribosomal protein S13 | B195_13486 | 99.72 | 96.36 | 95.8 | 96.36 | 96.36 | 96.92 |
| COG0100 | Ribosomal protein S11 | B195_13481 | 100 | 98.21 | 97.44 | 98.21 | 98.21 | 98.46 |
| **COG0102** | Ribosomal protein L13 | B195_12116 | 99.3 | 96.74 | 96.27 | 96.74 | 96.97 | 96.04 |
| **COG0103** | Ribosomal protein S9 | B195_12111 | 97.71 | 98.22 | 97.46 | 98.22 | 95.42 | 92.62 |
| **COG0124** | Histidyl-tRNA synthetase | B195_11566 | 90.47 | 88.84 | 90.54 | 88.84 | 87.52 | 86.98 |
| **COG0172** | Seryl-tRNA synthetase | B195_20565 | 91.18 | 90.94 | 85.71 | 90.94 | 88.6 | 86.34 |
| **COG0184** | Ribosomal protein S15P/S13E | B195_04417 | 97.78 | 94.81 | 95.19 | 94.81 | 90.41 | 94.82 |
| **COG0185** | Ribosomal protein S19 | B195_13576 | 98.91 | 98.91 | 98.91 | 98.91 | 97.46 | 97.46 |
| **COG0186** | Ribosomal protein S17 | B195_13551 | 97.38 | 96.63 | 97.0 | 96.63 | 96.63 | 97.00 |
| **COG0197** | Ribosomal protein L16/L10E | B195_13561 | 97.34 | 98.31 | 97.34 | 98.31 | 96.62 | 96.86 |
| **COG0200** | Ribosomal protein L15 | B195_13501 | 98.63 | 99.32 | 97.49 | 99.32 | 94.29 | 97.26 |
| **COG0201** | Preprotein translocase subunit SecY | B195_13496 | 97.97 | 97.89 | 98.04 | 97.89 | 96.61 | 96.92 |
| **COG0202** | DNA-directed RNA polymerase, alpha subunit | B195_13471 | 99.0 | 99.0 | 98.7 | 99.0 | 99.0 | 98.40 |
| **COG0215** | Cysteinyl-tRNA synthetase | B195_07500 | 89.95 | 89.3 | 87.93 | 89.3 | 84.82 | 88.01 |
| **COG0256** | Ribosomal protein L18 | B195_13516 | 98.58 | 99.15 | 97.72 | 99.15 | 97.44 | 97.15 |
| **COG0495** | Leucyl-tRNA synthetase | B195_13116 | 91.6 | 89.01 | 89.95 | 89.01 | 87.27 | 85.78 |
| **COG0522** | Ribosomal protein S4 and related proteins | B195_13476 | 98.71 | 98.71 | 98.23 | 98.71 | 97.91 | 98.23 |
| **COG0525** | Valyl-tRNA synthetase | B195_05521 | 94.2 | 93.33 | 93.4 | 93.33 | 91.29 | 89.78 |
| **COG0533** | Metal-dependent proteases with possible chaperone activity [UGMP family] | B195_13950 | 90.35 | 90.82 | 88.6 | 90.82 | 84.53 | 85.20 |
| **COG0541** | Signal recognition particle GTPase | B195_05675 | 92.08 | 90.99 | 91.29 | 90.99 | 90.86 | 89.33 |
| **COG0552** | Signal recognition particle GTPase [FtsY] | B195_14730 | 81.9 | 82.26 | 80.39 | 82.26 | 79.11 | 76.39 |
| **Average Identity** |  |  | **94.49** | **93.96** | **90.36** | **93.4** | **91.79** | **91.40** |

Therefore, we sought to use *in silico* tools for ANI analyses of genome sequence that are robust for identifying bacterial species if whole genome sequences are known. These tools were developed to establish not only the relationship between two genomes but also to substitute the experimentally derived DNA-DNA hybridization (DDH) method, a ‘gold standard’ for bacterial species identity (Goris *et al*., 2007). The JSpecies package (http://www.imedea.uib.es/jspecies) was among the first of its kind that was developed to calculate the ANI scores between a given pair of genomes for species delineation (Richter & Rossello-More, 2009). We compared the Lz4W^T^ genome similarity by JSpecies using selected *Pseudomonas* genomes that included the type strains of *P. fragi* species cluster and commonly studied *Pseudomonas* strains of environment (Table 4). In this analysis, ANIb values are derived from BLAST-N match data generated by pairwise comparison of artificially generated genomic segments of 1020 nucleotides that show more than 30% overall sequence identity over an alignable region of at least 70% of the 1020 bp length. ANIm values, on the other hand, are generated by MUMmer algorithm (the acronym “MUMmer” comes from “Maximal Unique Matches” used in the analysis) which can align sequences containing millions of nucleotide rapidly, using suffix trees (Richter & Rossello-More, 2009). JSpecies also gives output for the tetranucleotide frequency correlation coefficients (TETRA) which is an alignment-free parameter useful to phylogenetically short metagenome sequences. A threshold of 95-96% ANI score is recommended as putative boundary for species delineation, and ∼94% ANI value generally works well in mirroring the DDH range of ∼60-70%. We found that *P. deceptionensis* M1^T^ genome displays the highest similarity scores with the Lz4W^T^ genome, with ANIb and ANIm values being 85.25 and 87.02, respectively (Table 4). Other 5 type strains of the ‘*fragi*’ cluster (*P. fragi*, *P. psychrophila*, *P. taetrolens*, *P. lundensis*, and *P. endophytica*) also exhibited high ANI scores but below the species cut-off value. *P. fluorescens* Pf-01 genome displayed only 78.52 and 84.75 for ANIb and ANIm identity scores respectively. The TETRA values (0.97572, 0.96952, 0.96828, 0.97325, 0.93406, and 0.70533) of the six genomes of the ‘*fragi*’ cluster of species were consistent their relative ANI scores. Altogether, JSpecies analysis suggested that Lz4W^T^ genome identity does not cross the species threshold value (95%) when compared with the six closest type strains of the *P. fragi* cluster of species.

**Table 4.** JSpecies and GGDC based genome sequence analysis of Lz4W^T^ genome with related *Pseudomonas* species. The average nucleotide identity (ANIb and ANIm) scores and tetranucleotide frequency correlation coefficients (TETRA) were obtained by JSpecies, and DNA-DNA hybridization (DDH) values between the genomes were generated by GGDC.

| *Pseudomonas* species | Genome identity score with Lz4W<sup>T</sup> strain |  |  |  |
| --- | --- | --- | --- | --- |
|  | ANiB | ANIm | TETRA | DDH |
| *P. deceptionensis* DSM 26521<sup>T</sup> | 84.99 | 87.07 | 0.97572 | 29.9 |
| *P. fragi* B25<sup>T</sup> | 84.34 | 86.82 | 0.96951 | 29.3 |
| *P. psychrophila* DSM 17535<sup>T</sup> | 84.2 | 86.58 | 0.96829 | 28.8 |
| *P. taetrolens* DSM 21104<sup>T</sup> | 83.24 | 86.27 | 0.97324 | 27.8 |
| *P. helleri* DSM 29165<sup>T</sup> | 82.78 | 86.14 | 0.94073 | 27.4 |
| *P. weihenstephanensis* DSM 29166<sup>T</sup> | 82.11 | 85.82 | 0.899 | 26.4 |
| *P. lundensis* DSM 6252<sup>T</sup> | 81.78 | 85.72 | 0.93404 | 25.8 |
| *P. endophytica* BSTT44<sup>T</sup> | 80.99 | 85.18 | 0.70533 | 24.9 |
| *P. protegens* CHA0<sup>T</sup> | 78.62 | 84.84 | 0.89582 | 23.4 |
| *P. kilonensis* DSM13647<sup>T</sup> | 78.39 | 84.81 | 0.89486 | 23.7 |
| *P. extremaustralis* 14-3<sup>T</sup> | 78.29 | 84.95 | 0.88307 | 23.1 |
| *P. fluorescens* DSM 50090<sup>T</sup> | 78.05 | 84.92 | 0.8916 | 22.8 |
| *P. corrugata* NCPPB2445<sup>T</sup> | 77.94 | 84.71 | 0.91431 | 23.3 |
| *P. syringae* KCTC 12500<sup>T</sup> | 76.25 | 84.43 | 0.93187 | 22.3 |
| *P. putida* NBRC 14164<sup>T</sup> | 75.63 | 83.83 | 0.89719 | 21.7 |
| *P. guinea* LMG24016<sup>T</sup> | 73.88 | 83.83 | 0.79678 | 20.4 |
| *P. aeruginosa* DSM 50071<sup>T</sup> | 73.53 | 83.09 | 0.79556 | 20.3 |
| *P. protegens* Pf-5 | 78.63 | 84.82 | 0.89564 | 23.4 |
| *P. fluorescens* Pf0-1 | 78.28 | 84.87 | 0.89653 | 23.1 |
| *P. fluorescens* SBW25 | 78.07 | 84.91 | 0.86612 | 23.1 |
| *P. syringae* pv. *tomato* DC3000 | 76.27 | 84.39 | 0.92754 | 22.1 |

We also performed Genome Blast Distance Phylogeny (GBDP) analysis wherein genome to genome distances of different bacteria are inferred using HSPs (high-scoring segment pairs) matches of the genomes, and DDH values are calculated from the intergenomic distance using specific distance formula (Meier-Kolthoff *et al*., 2013). The GBDP predicted DDH values of Lz4W^T^ genome are shown along with ANI identity scores in Table 4. The values were obtained using the formula 2 of GGDC 2.0 (GBDP2_BLASTPLUS program) software as recommended (http://ggdc.dsmz.de) for comparing draft genomes. GGDC compares a query genome with reference genome to calculate the intergenomic distance under three different distance formulae, of which Formula 2 is independent of the genome lengths and is thus robust against the use of incomplete draft genomes. In our analysis of Lz4W^T^ genome against the genome sequences of *P. deceptionensis* M1^T^, *P. fragi* ATCC4973^T^, *P. psychrophila* E-3^T^, *P. lundensis* DSM6252^T^, and *P. endophytica* BSTT44^T^ the predicted DDH values were 29.9, 29.2, 28.8, 25.8, and 25.0%, respectively. These values are quite low, as the probability that DDH values >70% (i.e., same species) between the Lz4W^T^ and six type strains of “*fragi*” cluster of genomes were determined to be only 0.07% (via logistic regression) as per the Formula 2. Thus, GBDP analysis also suggests that Lz4W^T^ is a novel species of *Pseudomonas*.

To summarize, Lz4W^T^ strain clearly differs from the major phenotypic characteristics of “*syringae*” species (Table 1). The five major type strains of the “*fragi*” cluster of species (*P. deceptionensis* M1^T^, *P. endophytica* BSTT44^T^, *P. fragi* ATCC4973^T^, *P. psychrophila* E-3^T^, and *P. lundensis* DSM6252^T^) which exhibit very high 16S rRNA gene sequence identity (99.72 to 99.45%) with Lz4W^T^ (Table S1) can be differentiated from each other by genomic criteria (Table 4) and by phenotypic characteristics (Table 5). The most closely related genomes of three type strains (*P. deceptionensis* M1^T^, *P. fragi* ATCC4973^T^, and *P. psychrophila* E-3^T^) and Lz4W^T^ displayed lower than the required species cutoff values by SpecI (98.5%) and JSpecies ANI scores (95%). More importantly, predicted DDH values (29.9 % or below) of Lz4W^T^ with the above type strains falls far below the species cut-off value (>70%). Hence, we believe Lz4W^T^ is a novel species of *Pseudomonas*, for which we propose the name *Pseudomonas cryophila* sp. nov. (cry.o’phi.la. Gr. n. *kryos*, icy cold, frost; N.L. adj. *philus* –*a* –*um* (from Gr. adj. *philos* –*ê* – *on*), friend, loving; N.L. fem. adj. *cryophila*, cold-loving).

**Table 5.**
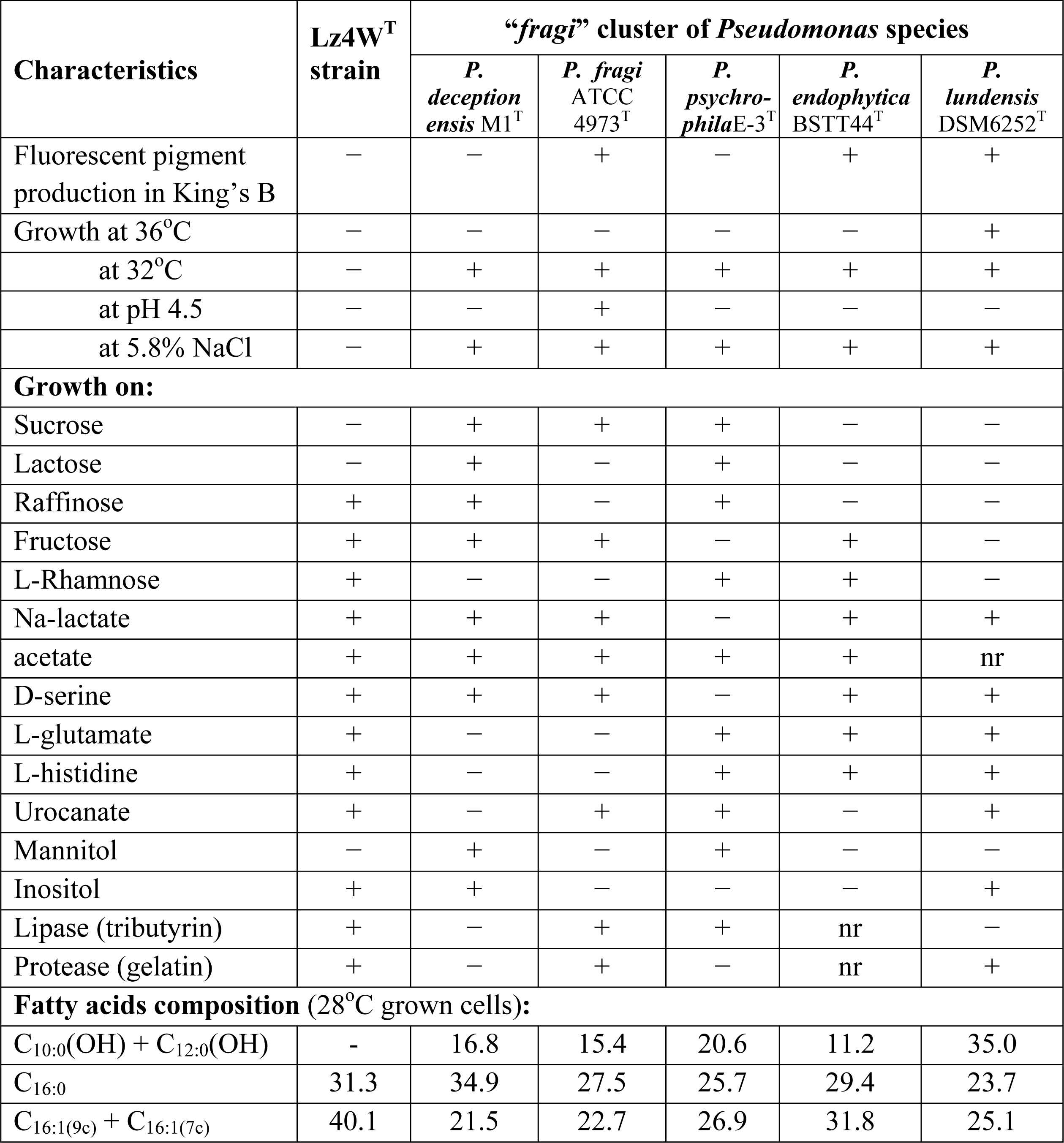
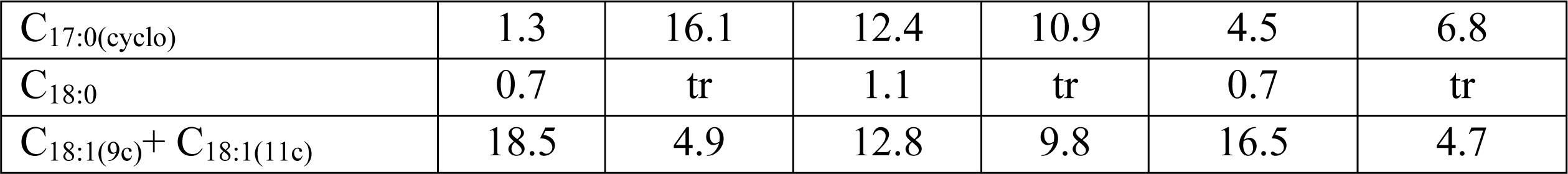
Differential phenotypic characteristics of Lz4W^T^ and the type strains of phylogenetically closest *Pseudomonas* species of “*fragi*” cluster. The data have been compiled from Carrion *et al* (2011), Ramirez-Bahena *et al* (2015), Yumoto *et al* (2001), Kiran *et al* (2004), Shivaji *et al* (1989) and this study. A ‘+’ and ‘−’ indicates positive and negative for the tests, respectively; nr, not reported.

Interestingly, we note that the draft genome sequences of two putative “*fragi*” strains, *P. fragi* A22 (acc no. AHZY01000000) and *P. fragi* P121 (acc no. CP013861) reported by Mei *et al* (2012) and Yanzhen *et al* (2016) respectively, are almost identical to Lz4W^T^ genome sequence. The genome similarity criteria including the ANI scores by JSpecies and DNA-DNA hybridization (DDH) index by GGDC method (Table 6) show that the “*fragi*” isolates A22 and P121 are closest to *P. cryophila* Lz4W^T^, and they are as much diverged as Lz4W^T^ from the *P. fragi* ATCC4973^T^ type strain. The high DDH values (above 99.4%) between the three strains (Lz4W^T^, “*fragi* A22”, and “*fragi* P121”) put them apart from the *P. fragi* ATCC4973^T^ type strain. Therefore, we suggest that *P. fragi* strains A22 and P121 should also be classified as *P. cryophila* species. Interestingly, the strains A121 and P22 were isolated from the Arctic soil and water respectively (Yanzhen *et al*. 2016, Mei *et al*. 2012), which adds a new dimension to the origin and geographical distribution of *P. cryophila* species in two opposite poles of our globe.

**Table 6.** Genome identity (ANIb, ANIm, TETRA, and DDH) scores of *P. cryophila* Lz4W^T^ with “*P. fragi* A22” and “*P. fragi* P121” strains, which distance them from the *P. fragi* ATCC4973^T^ type strain and other ‘*fragi*’ cluster of related species (*P. deceptionensis* M1^T^, *P. psychrophila* E-3^T^, *P. lundensis* DSM6252^T^, *P. endophytica* BSTT44^T^, and *P. taetrolens* DSM21104^T^) that exhibited very high 16S rRNA gene homology.

| *Pseudomonas*<br>species | Genome Accession<br>no. | Genome<br>Size (bp) | GC<br>percent | Identity score with *P. cryophila* Lz4W<sup>T</sup> |  |  |  |  |
| --- | --- | --- | --- | --- | --- | --- | --- | --- |
|  |  |  |  | 16S<br>rRNA | ANiB | ANIm | TETRA | DDH |
| *P. cryophila* Lz4W<sup>T</sup> | AOGS000000000 | 4982906 | 58.6 | **100** | **100** | **100** | **1.00000** | **100** |
| *P. fragi* A22 | AHZY01000000 | 5043229 | 58.6 | **98.76** | **99.2** | **99.35** | **0.99986** | **94.8** |
| *P. fragi* P121 | CP013861 | 5101809 | 58.6 | **99.93** | **99.21** | **99.29** | **0.99972** | **94.4** |
| *P. fragi* ATCC4973<sup>T</sup> | AHZX01000000 | 5000569 | 59.3 | 99.45 | 83.86 | 86.82 | 0.96951 | 29.2 |
| *P. deceptionensis*<br>M1<sup>T</sup> | JYKX01000000 | 5052933 | 58.6 | 99.72 | 84.63 | 87.07 | 0.97572 | 29.9 |
| *P. psychrophila* E-3<sup>T</sup> | JYKZ00000000 | 5334010 | 57.5 | 99.46 | 83.70 | 86.58 | 0.96829 | 28.8 |
| *P. lundensis*<br>DSM6252<sup>T</sup> | JYKY00000000.1 | 4992500 | 58.5 | 99.46 | 82.22 | 85.68 | 0.93404 | 25.8 |
| *P. endophytica*<br>BSTT44<sup>T</sup> | NZ_<br>LLWH00000000.1 | 4973838 | 55.2 | 99.52 | 81.02 | 85.18 | 0.70533 | 25.0 |
| *P. taetrolens*<br>DSM21104<sup>T</sup> | JYLA00000000.1 | 4921904 | 58.3 | 98.73 | 83.61 | 86.21 | 0.97325 | 27.8 |

Description of *Pseudomonas cryophila* sp. nov.

*Pseudomonas cryophila* (cry.o’phi.la. Gr. n. *kryos*, icy cold, frost; N.L. adj. *philus* –*a* –*um* (from Gr. adj. *philos* –*ê* –*on*), friend, loving; N.L. fem. adj. *cryophila*, cold-loving).

Basonym: *Pseudomonas syringae* Lz4W (Shivaji *et al*., 1989).

The description of this taxon is the same as that given by Reddy et al (1989) for *Pseudomonas syringae* Lz4W, except for few key revised characteristics that have been specified here. The species is Gram-negative, aerobic, motile rod-shaped cells with one polar flagellum, and displays temperature and nutrient dependent cell-size variation, being smaller at low-temperature and in low-nutrients. Growth temperature ranges between 0 – 30°C with optimum growth at 22°C. Grows best without NaCl and does not grow in 5.8% NaCl. Grows in a pH range of 5.0 to 9.0 with optimum at 7.0. Tests positive for oxidase, catalase, arginine dihydrolase, and urease but negative for β-galactosidase, esculin hydrolase, indole production, nitrate reduction. Positive for phosphatase, gelatinase, and lipase activity on tributyrin but not on Tween 80. Does not produce fluorescent pigment in King’s B medium, and does not show hyper-sensitive reaction in tobacco. Tests positive for assimilation of pyruvate, citrate, succinate, fumarate, α-ketoglutarate, acetate, lactate, D-glucouronate, D-tartarate, raffinose, fructose, L-rhamnose, glycerol, and inositol. Tests negative for assimilation of sucrose and mannitol, and no levan formation. Tests positive for utilization of all naturally occurring L-amino acids as carbon and nitrogen sources, except for three (methionine, cysteine, and tryptophan). The major fatty acids in 22°C grown cells are C_16:0_ (28.9%), C_16:1(9c)_ (41.7%), and C_18:1(11c)_ (18.5%) and low C_17:0_ cyclo (1.4%).

The type strain is Lz4W^T^ (=CFBP8403^T^ =KCTC42933^T^ =LMG29591^T^ =MTCC 673^T^). The genomic G + C content of the type strain is 58.7 mol%.

## Abbreviations

ANI: average nucleotide identity
DDH: DNA-DNA hybridization
PMGs: phylogenetic marker genes
SpecI: species identification tool
GBDP: Genome Blast Distance phylogeny

## Acknowledgements

We are grateful to Dr. Cindy E. Morris (INRA, Avignon, France) for her correspondence on the Lz4W species identity and verifying some of the characteristics of the bacterial strain. We thank Dr. G.S.N. Reddy (CCMB, Hyderabad) for re-assessing the gelatinase and lipase activities of Lz4W strain. This work was supported by the CSIR (Council of Scientific and Industrial Research) Grants under GENESIS (BSC0121) and EpiHeD (BSC0118) projects to MKR. AP is also supported by a senior research fellowship from the CSIR.

